# Comparative Phylogenomic Synteny Network Analysis of Mammalian and Angiosperm Genomes

**DOI:** 10.1101/246736

**Authors:** Tao Zhao, M. Eric Schranz

**Affiliations:** Biosystematics Group, Wageningen University, 6708 PB Wageningen, The Netherlands

**Keywords:** synteny, networks, genome evolution, gene family dynamics, phylogenetic profiling, mammals, angiosperms

## Abstract

**Background:** Synteny analysis is a valuable approach for understanding eukaryotic gene and genome evolution, but still relies largely on pairwise or reference-based comparisons. Network approaches can be utilized to expand large-scale phylogenomic microsynteny studies. There is now a wealth of completed mammalian (animal) and angiosperm (plant) genomes, two very important lineages that have evolved and radiated over the last ~170 million years. Genomic organization and conservation differs greatly between these two groups; however, a systematic and comparative characterization of synteny between the two lineages using the same approaches and metrics has not been undertaken.

**Results:** We have built complete microsynteny networks for 87 mammalian and 107 angiosperm genomes, which contain 1,464,753 nodes (genes) and 49,426,268 edges (syntenic connections between genes) for mammals, and 2,234,461 nodes and 46,938,272 edges for angiosperms, respectively. Exploiting network statistics, we present the functional characteristics of extremely conserved and diversified gene families. We summarize the features of all syntenic gene clusters and present lineage-wide phylogenetic profiling, revealing intriguing sub-clade lineage-specific clusters. We depict several representative clusters of important developmental genes in humans, such as *CENPJ, p53* and *NFE2*. Finally, we present the complete homeobox gene family networks for both mammals (including Hox and ParaHox gene clusters) and angiosperms.

**Conclusions:** Our results illustrate and quantify overall synteny conservation and diversification properties of all annotated genes for mammals and angiosperms and show that plant genomes are in general more dynamic.

## Background

The patterns and differences of gene and genome duplication, gene loss, gene transpositions and chromosomal rearrangements can inform how genes and gene families have evolved to regulate and generate (and potentially constrain) the amazing biological diversity on Earth today. For comparative genomics, synteny reflects important relationships between the genomic context of genes both in terms of function and regulation and is often used as a proxy for the constraint and/or conservation of gene function [1, 2]. Thus, syntenic relationships across a wide range of species provide crucial information to address fundamental questions on the evolution of gene families that regulate important traits. Synteny data can also be very valuable for assessing and assigning gene orthology relationships, particularly for large multigene families where phylogenetic methods maybe non-conclusive [1, 3, 4]. Synteny was originally defined as pairs or sets of genes located on homologous chromosomes in two or more species, but not necessarily in the same order [5]. However, the current widespread usage of the term synteny, which we adopt, implies conserved collinearity and genomic context.

While the basic tenants of gene and genome organization and evolution are similar across major eukaryote lineages, there are also significant differences that are not fully characterized nor understood. For example, the length and complexity of genes and promoters, the types of gene families (shared or lineage-specific), transposon density, higher-order chromatin domains and the organization of chromosomes can differ significantly between plants, animals and other eukaryotes [6–9]. In general, genome organization and gene collinearity is substantially less conserved in plants than in mammals. One major characteristic of flowering plant genomes is the prevalent signature of shared and/or lineage-specific whole genome duplications (WGDs) [10–15]. While the genomes of mammalian vertebrates show evidence of only two shared and very old rounds of WGD; often referred to as “2R” [16–18]. The variation in genomic organization between lineages is partially due to differences in fundamental molecular processes such as DNA-repair and recombination, but also likely reflect the historical biology of groups (such as mode of reproduction, generation times and relative population sizes). Differences in gene family and genome dynamics have significant effects on our ability to detect and analyze synteny.

While the number of quality reference genomes is growing exponentially, a major challenge is how to detect, represent, and visualize synteny relations of all members from a gene family across many genomes simultaneously. Conventional dot plots display macroscale collinear blocks between/within only two genomes in two-dimensional images. Parallel coordinate plots (like CoGe SynFind [19, 20]) describe collinear blocks surrounding a locus identifier and visualize the blocks at the local genomic scale. With the abundance of new genomic data, the changes for multispecies collinearity visualization are only exacerbated. We have developed a network-based approach to organize and display local synteny [21, 22] and have applied it to understand the evolution of the entire MADS-box transcription factor family across 51 plant genomes as a proof of principle of the method [22]. We identified several evolutionary patterns including extensive pan-angiosperm retention of certain gene clades, ancient retained tandem duplications and lineage-specific transpositions such as the floral patterning genes in Brassicaceae [22]. Our approach can be scaled to analyze not just one gene family, but all gene families across a lineage.

The aim of this study is to investigate and compare the dynamics and properties of the entire synteny networks of all annotated genes for mammals and angiosperms. To this end, we analyzed the syntenic properties of 87 mammalian and 107 plant genomes (Figure 1) which represent most major phylogenetic clades of both mammalian and angiosperm groups across ~170 million years of evolution [13, 23–25]. For mammals, the species used covered the three main clades of Afrotheria, Euarchontoglires, and Laurasiatheria, as well as first-branching groups like *Ornithorhynchus anatinus* (platypus). For angiosperms, the species also cover three main groups of Monocots, Superasterids, and Rosids, as well as first-branching groups such as *Amborella trichopoda* (Figure 1). Some clades are more heavily represented than others such as primates (human relatives) and crucifers (Arabidopsis relatives) due to research sampling biases. Regardless, most major lineages are represented. Also, there are differences in the overall quality and completeness of the genome assemblies used, but this was a factor we wanted to analyze and assess using synteny analysis.

**Figure 1.**
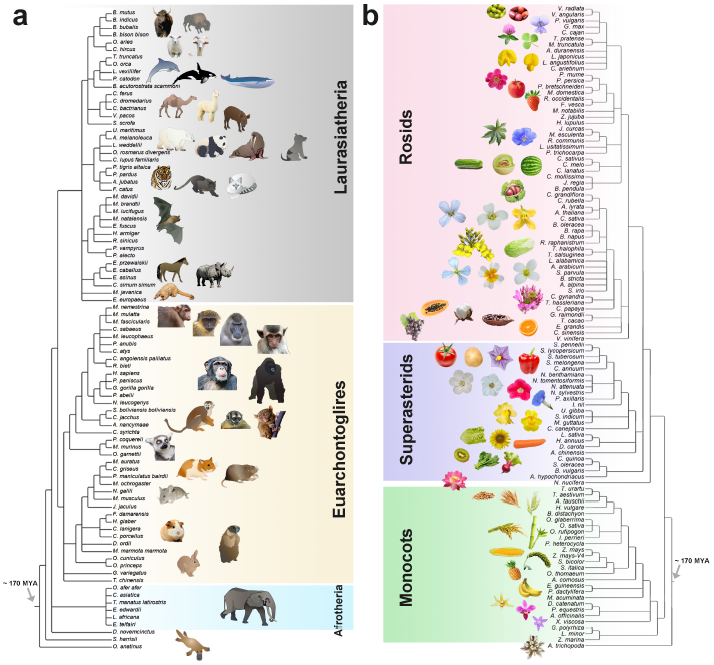
Phylogenetic relationships of mammal and angiosperm genomes analyzed. (a) Mammal genomes used, highlighting the three main placental clades Afrotheria, Euarchontoglires and Laurasiatherias. (b) Angiosperm genomes used, highlighting the three main clades Monocots, Superasterids and Rosids.

## Results and discussion

### Genome collection, pairwise synteny comparisons

We used fully-sequenced genomes to investigate all syntenic blocks within and across genomes. Initially we searched public databases maintaining mammalian and angiosperms genome resources such as NCBI, Ensembl, CoGe and Phytozome. Candidate genomes had to contain downloadable complete predicted gene models and gene position annotations. Ultimately, we analyzed 87 mammalian genomes, presented according to the consensus species tree adopted from NCBI taxonomy (Figure 1, Supplemental Table 1) which included 1 Prototheia (*Ornithorhynchus anatinus*), 1 Metatheria (*Sarcophilus harrisii*), 1 Xenarthra (*Dasypus novemcinctus*), 6 Afrotheria, 38 Euarchontoglires and 40 Laurasiatheria species. For angiosperms, we analyzed 107 genomes including 1 Amborellaceae (*Amborella trichopoda*), 26 Monocots (including 14 Poaceae) and 80 eudicots (including 1 Proteales (*Nelumbo nucifera*), 23 Superasterids (Asterids and Caryophyllales), and 56 Rosids) (Figure 1, Supplemental Table 1).

We modified all peptide sequence files and genome annotation GFF/BED files with corresponding species abbreviation identifiers, followed by pairwise all-vs-all genome comparisons for synteny block detection [as described in 21, 22]. To assess the overall impact of phylogenetic distance, genome assembly quality and/or genome complexity, we summarized the number of syntenic gene pairs for all pairwise genome comparisons (7,569 times for mammals and 11,449 times for angiosperms) into color-scaled matrixes (Figure 2) organized using the same species phylogenetic order as in Figure 1.

**Figure 2.**
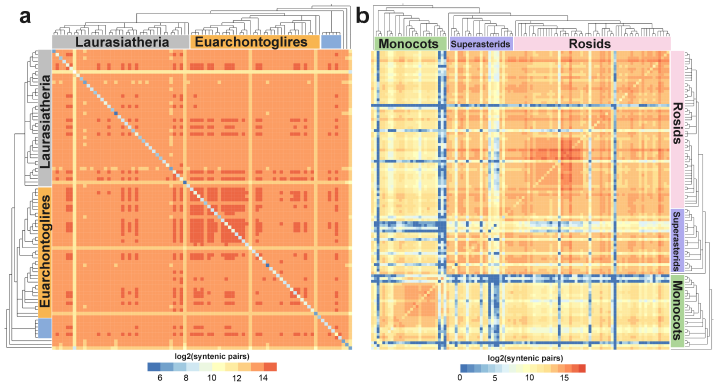
Pairwise synteny comparisons of mammal and angiosperm genomes. (a)Pairwise synteny comparison across Mammal genomes. (b) Pairwise synteny comparison across Angiosperm genomes. The logarithmic color-scale indicates the number of syntenic gene pairs. Species are ordered according to the consensus phylogeny (Figure 1). Overall, average synteny is much higher across mammals than plants. Also, there is a stronger phylogenetic signal seen for plant genomes. The method also allows for easy detection of potentially low-quality genomes (overall lower syntenic pair scores). The diagonal for both plots represents intra-genome comparisons which can detect potential recent and ancient WGDs. Note, that almost all plant genomes have higher intra-genome syntenic pair scores than all mammal intra-genome comparisons.

The diagonal of the matrix represents self- vs. self-contrasts and indicates the number of retained duplicate genes, which is indicative of recent and/or ancient WGDs. The lighter orange and blue rows with fewer syntenic links could reflect key biological or genomic differences, but is much more likely to be due to poor quality genome assemblies. For example, the mammalian genomes of *O. anatinus*, *Galeopterus variegatus*, *Carlito syrichta, Manis javanica*, and *Tursiops truncates* (Figure 2a) and for angiosperms *Humulus lupulus*, *Triticum urartu*, *Aegilops tauschii*, and *Lemna minor* (Figure 2b).

As shown in the matrixes, mammalian genomes overall are in general highly syntenic regardless of phylogenetic distance (Figure 2a) with primate vs primate comparisons showing marginally higher scores. Whereas plant genomes show more phylogenetic signal (e.g. monocots vs monocots and crucifers vs. crucifers), the impact of recent WGD (e.g. *Brassica napus*) and more variability overall (due to assemblies from different groups of researchers, different qualities, multiple independent WGDs) (Figure 2b). Note, that almost all plant genomes have higher intra-genome syntenic pair scores than all mammal intra-genome comparisons. We further checked genome characters by plotting syntenic gene percentage against Pfam annotation percentage for each genome (Supplemental Figure 1). Based on these results, we removed four poor-quality plant genomes (*H.lupulus*, *T. urartu*, *A. tauschii*, and *L. minor*) before proceeding to the next step of our analyses.

### Characterization of synteny networks

The entire synteny networks are composed of all syntenic genes identified within all the syntenic blocks. Specifically, there are 1,464,753 nodes (genes) and 49,426,268 edges (syntenic connections between genes) for mammals, and 2,234,461 nodes and 46,938,272 edges for angiosperms, respectively. To evaluate genomic conservation of gene families (for gene family assignments see Methods) over evolutionary time scales from the synteny network data, we introduce two estimators: average clustering coefficient (Supplementary Figure 2) and the percentage of genes in the family that are syntenic (syntenic percentage) for every gene-family (Figure 3a). A clustering coefficient is calculated for all nodes in the synteny network, as a measure of the degree to which nodes in a graph tend to cluster together. Genes can be mobilized (e.g. transposed) to other genomic contexts (e.g. unique or lineage-specific contexts) and thus will no longer be collinear or syntenic to other species or lineages. Thus, we use percentage (gene family members in the network/ total gene family members in the genomes) to quantify the proportion of the genes retaining synteny.

**Figure 3.**
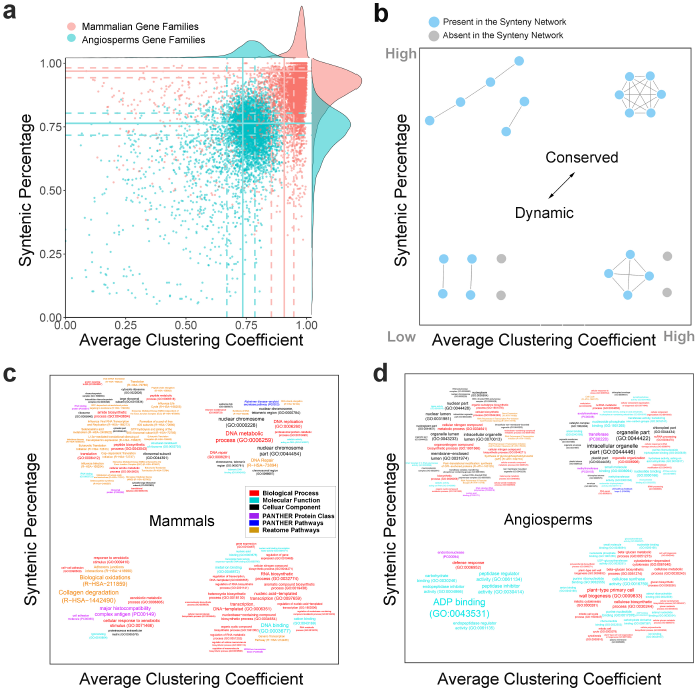
Network properties of gene families from mammal and angiosperm genomes. (a) Distributions of gene family dynamics of mammal (11,830 in red) and angiosperm (10,617 in blue) gene families plotted using percentage of syntenic genes and average clustering coefficients per family. Quartiles of average clustering coefficient and syntenic percentage for both mammals and angiosperms are indicated by dashed (25%/75%) and solid (median) lines. (b) Conceptual model depicting different patterns of synteny network connectivity, according to data distribution, with further analysis based on 25% quartiles. (c, d) Comparative word clouds based on upper and lower quartiles for functional enrichment of significant terms with representative C-P profiles for mammals (c) and angiosperms (d). Font sizes are representative of adjusted p-values.

We then plotted the average clustering coefficient and retention percentage of all the gene families for the mammalian (11,830 gene families) and angiosperm (10,617 gene families) synteny networks (Figure 3a). Mammalian gene families overall have significantly higher clustering coefficients (mean 0.92 for mammals compared to 0.72 for angiosperms; *P* < 0.001, Wilcoxon-Matt-Whitney test) and retention percentage (mean 0.88 for mammals compared to 0.71 for angiosperm; *P* < 0.001, Wilcoxon-Matt-Whitney test) than that of angiosperms (Figure 3a). This confirms that over large evolutionary time scales, genomic context is generally more conserved and constrained in mammals than for angiosperms.

Syntenic dynamics of all gene families could be classified and compared to other gene families by our C-P (Clustering coefficient vs Percentage) quartile analysis method, as conceptually depicted in Figure 3b. We defined values of the top 25% quartile as “high”, and the bottom 25% quartile as “low” for both mammals and angiosperms. The resulting four categories are highlighted (Figure 3b). The high clustering coefficient plus high retention percentage in the synteny network (“high-high” C-P values), indicates the both most syntenically conserved and most completely syntenic gene families, and thus the most inter-connected networks (Figure 3b, Supplementary Table 2). Genes in the category of “high-low” C-P detect gene families where certain gene sub-families and/or phylogenetic clades are highly syntenic, but overall many gene members are absent from the clusters (thus a low percentage). Non-syntenically connected gene family members may be prone to transposition (Figure 3b, Supplementary Table 2). In contrast, the category “low-high” C-P means that a high proportion of the gene family members are in the network, but not always well connected, for example due to tandem gene cluster expansions (Figure 3b, Supplementary Table 2). Lastly, the category “low-low” C-P represent gene families that are distributed dispersedly (such as across pericentromeric regions) and thus non-syntenic, or represent young transpositions or lineage-specific genes shared only between a small number or related species (Figure 3b, Supplementary Table 2).

### Comparative synteny dynamics of gene families of mammals and angiosperms

We investigated if gene families with similar C-P synteny dynamics (high-high, high-low, low-high, and low-low), might also have similar functional annotations (e.g. GO terms) [26, 27]. We tested for pathway and gene-function enrichment of gene families within each of the four C-P profiles for both mammals and angiosperms (Figure 3c and 3d). Over-representative terms are shown in a word-cloud with font sizes indicating the p-value (Fisher’s exact test with Bonferroni correction). For mammals, gene families with “high-high” profiles are functionally enriched in DNA metabolic processes, such as “DNA replication” and “DNA repair”. Interestingly Alzheimer disease-amyloid secretase pathway (P00003) genes are enriched in this category (Figure 3c). By contrast, “low-low” gene families include functions in immune responses and pathways (e.g., “cellular response to xenobiotic stimulus”, “Collagen degradation”, “Biological oxidations”), enriched protein classes are “major histocompatibility complex antigen (PC00149)” and “cell adhesion molecule (PC00069)” (Figure 3c). The mammalian “high-low” group is enriched for genes that function in DNA-templated gene transcription and DNA binding, such as KRAB box transcription factors (PC00029) [28] (Figure 3c). As transcription factors bind specific promoters and thus regulate a variety of developmental and environmental processes. Moreover, transcription factors commonly consist of multiple members. Thus, it can be hypothesized that some gene family members are highly conserved and genomically constrained, while other members are versatile and transposed into new genomic positions. Finally the “low-high” group is enriched for genes involved in translation (e.g. “peptide biosynthetic process”, “peptide metabolic process”) and ribosomal component (e.g. “ribosomal subunit”, “ribonucleoprotein complex”), most enriched Reactome Pathways are closely related to translation processes (e.g. “eukaryotic translation”, “Cap-dependent translation initiation”), as well as infectious disease related pathways (e.g. “Influenza infection”, “Influenza life cycle”, and “Influenza viral RNA transcription and replication”) (Figure 3c).

The functional enrichment analysis of angiosperms shows a different pattern than for mammals (Figure 3d). Plant “high-high” gene families are enriched for organelle components (e.g. “organelle part”, “intracellular organelle”, “chloroplast part”, “organelle organization”, and “plastid part”), as well as acetyltransferase, transferase and methyltransferase proteins for the processes such as “DNA repair”, “ncRNA metabolic process” and “methylation” (Figure 3d). Many of these categories are plant-specific related to photosynthesis. By contrast, the plant “low-low” group is enriched by defense response genes such as “peptidase inhibitor activity”, “endopeptidase inhibitor”, and “ADP binding”. “Low-high” gene families function in nuclear part components (e.g. “intracellular organelle lumen”, “organelle lumen”), biosynthetic process (e.g. “organonitrogen compound biosynthetic process”, “cellular aromatic compound metabolic process”), cell surface proteins (e.g. “synthesis of glycosylphosphatidylinositol (GPI)) and gene expression (e.g. “RNA polymerase complex”, “nucleic acid binding”, “RNA polymerase II transcription initiation”). Interestingly, “high-low” part of plant genes function in cell wall (e.g. “plant-type primary cell wall biogenesis”, “cellulose biosynthetic process”, “beta-glucan biosynthetic process”) (Figure 3d). Classifying and characterizing gene families according to their “synteny network C-P” scores allows for the relative comparisons of any gene family to all others across a lineage (Supplementary Table 2). The degree of conservation likely reflects functional constraints of the family. For example, gene families with a high-high C-P are responsible for fundamental functions (i.e. DNA repair and photosynthesis.) and low-low C-P gene families are highly mobile and functionally flexible (such as both animal and plant NLR family defense-related receptors [29] and plant P450s and F-box genes) (Supplementary Table 2).

### Comparative synteny network clustering

We next performed a clustering analysis for the entire mammal and angiosperm synteny networks. We used Infomap [30] as the clustering algorithm due to its efficiency and accuracy in handling large graphs with millions of nodes and because it has consistently out-performed other available methods [31]. The clustering results for mammals and angiosperms are summarized and compared in terms of cluster-size distributions (Figure 4a and 4b), corresponding clustering coefficients (Figure 4c and 4d), and number of species included per cluster (Figure 4e and 4f).

**Figure 4.**
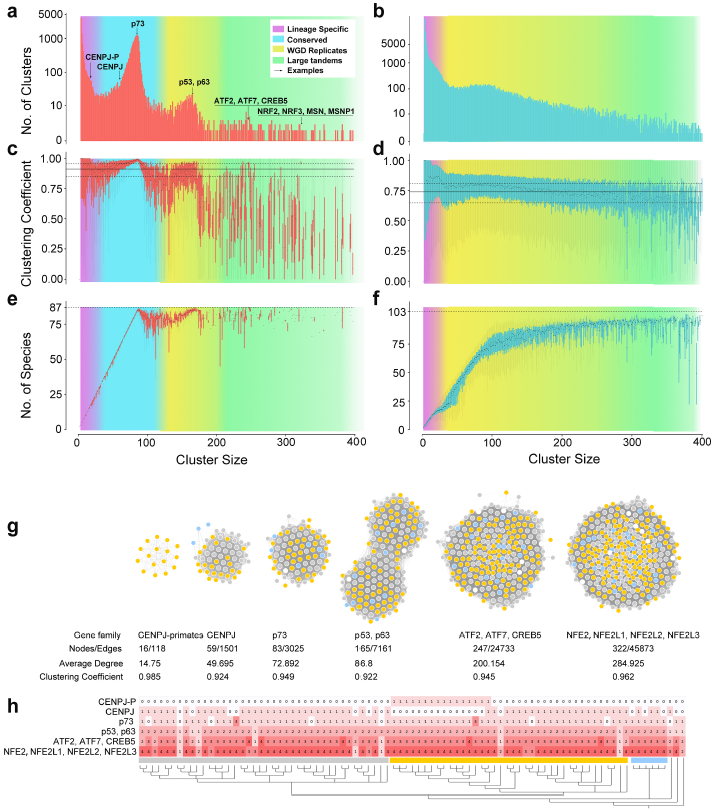
Synteny cluster statistics of mammal and angiosperm genomes and representative mammalian synteny clusters. Approximate size ranges for clusters oflineage-specific, conserved, WGD replicates, and large tandem genes are shaded in purple, cyan, yellow, and olive green, respectively. (a) Sizes distribution of all mammalian gene syntenic clusters. Representative examples are pointed and labeled on the curve. (b) Sizes distribution of all angiosperms gene syntenic clusters (c) Boxplot of clustering coefficient by mammalian cluster sizes. (d) Boxplot of clustering coefficient by angiosperm cluster sizes. (e) Number of involving genomes for mammalian clusters by cluster sizes. Number of involving genomes for angiosperm clusters by cluster sizes. (g) Six representative and diverse mammalian clusters of CENPJ (primate-specific one and the others), p73, p53-p63, ATF2-ATF7-CREB5, and NFE2-NFE2L1-NFE2L2-NFE2L3. Total number of nodes, edges, average degree, and clustering coefficient are indicated accordingly below. (h) Phylogenetic profiling of the clusters from (g), a color gradient of red indicates the number of syntelogs in each species.

Mammalian genomes have a prevalent peak of syntenic gene families that are present only once per taxa (single copy orthologous gene cluster peak shaded in cyan, Figure 4a). To the right, there is a second modest peak of duplicated (ohnolog) genes due to the ancient 2R WGD events (shaded in bright yellow, Figure 4a). These two peaks could be further explained by Figure 4c and Figure 4e that depict the corresponding average clustering coefficient and number of species, respectively. We observe that the peak in cyan in Figure 4a is accompanied by a steady increasing trend of the clustering coefficient and the number of species involved (Figure 4c). A similar trend was observed for the clusters forming the peak in yellow due to WGD (Fig 4a). On the far left there is the rather modest proportion of lineage specific genes (clusters of syntenic genes between only a subset of mammalian species or clade(s) (shaded in purple, Figure 4a). On the far right are large multigene clusters usually with multiple syntenic gene copies conserved across multiple species due to tandem duplications such the well-known Hox-genes (shaded in olive green, Figure 4a). Representative examples are labeled on the curve, and further depicted in Figure 4g and Figure 4h.

In contrast, angiosperm genomes show a very large proportion of lineage-specific clusters on the far left (shaded in purple, Figure 4b). The clustering coefficients for these clusters is often above the threshold of “high” (top 25%, which was defined earlier for the C-P classification) (Figure 4d) and the cluster size for these lineage-specific clusters is mostly between 10 to 30 (shaded in cyan, Figure 4f), reflecting the number of species and gene copies within particular phylogenetic groups such as Fabaceae, Brassicaceae, and Poaceae. Next, a rather broad peak of gene clusters are observed that are conserved across many lineages (Figure 4b) of genes that are single-copy in some lineages and in two/more copies in other lineages due to WGD. Also, there is a larger proportion of large multigene families seen to the far right (shaded in olive green, Figure 4b). There is a variation for the number of species per cluster for these large multi-gene families in angiosperms (Figure 4f).

The combination of cluster size, corresponding clustering coefficient, and number of involved species were used to select representative synteny clusters for mammals. As an example of a lineage-specific cluster we show *CENPJ* (as an example an of a primate lineage-specific cluster), *p73* as an example of a single copy conserved cluster, *p53-p63* as an example of 2-ohnologs-retained WGD cluster, *ATF2-ATF7-CREB5* as an example of 3-ohnolog-retained WGD cluster, and *NFE2-NFE2L1-NFE2L2-NFE2L3* as example of 4-ohnolog-retained WGD cluster (Figure 4a, 4g and 4h). It has been reported that *CENPJ* regulates brain size [32, 33], and primates have relatively larger brains [34, 35]. It is interesting that we found primates formed a lineage-specific *CENPJ* synteny cluster (Figure 4g and 4h) compared to other mammals. This indicates that *CENPJ* underwent a gene transposition event at or near the divergence of the primate ancestor from other mammals. Thus, the primate gene copy is in a unique genomic context facilitating potential new/altered regulatory patterns and gene functions. The *p53, p63* and *p73* genes compose a family of transcription factors involved in cell response to stress and development [36, 37]. *p63* is previously perceived close related to *p73* because of the similar protein domain compositions, however our result shows *p63* and *p53* are ohnolog duplicates retained after WGD. Other ohnolog clusters with strong support from our analyses include *ATF2-ATF7-CREB5*, transcription factors with broad roles such as activating CRE-dependent transcription, cancer progression and immunological memory [38–41] and *NFE2-NFE2L1-NFE2L2-NFE2L3*, also with broad roles such as regulation of oxidative stress, aging and cancer cell proliferation [42–44].

### Comparative phylogenetic profiling of synteny clusters

To further visualize and understand genomic diversity, we performed phylogenetic profiling of all synteny clusters of mammals and angiosperms (Figure 5a and 5b). Blue columns indicate conserved single copy syntenic clusters, orange columns indicate retained duplicate copy clusters (i.e. conserved ohnologs from WGD), and the red columns signify conserved clusters with more than two copies (e.g. conserved tandem clusters) (Figure 5a and 5b). Nearly empty rows of the less-syntenic species are consistent with the pairwise matrix in Figure 2.

**Figure 5.**
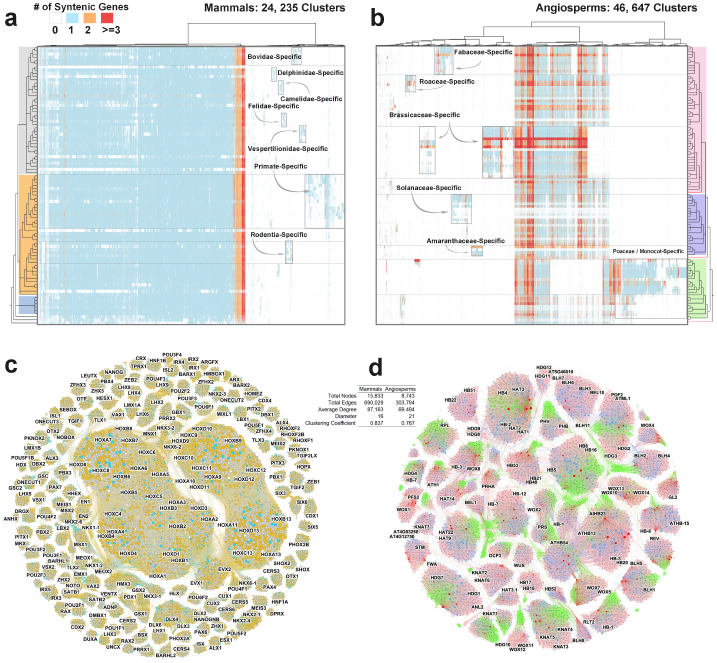
Phylogenetic profiling of all synteny clusters and complete Homeodomain multigene family synteny networks from mammal and angiosperm genomes. (a) Phylogenetic profiling of all synteny clusters and complete Homeodomain multigene family synteny networks from mammal and angiosperm genomes. (a) Phylogenetic profiling of all mammalian clusters (size >= 2). Groups of lineage-specific clusters are boxed and labeled. (b) Phylogenetic profiling of all angiosperm clusters (size >= 3). Groups of lineage-specific clusters are boxed and labeled. (c, d) Synteny network of all homeo-domain proteins for mammals (c) and angiosperms (d), representative *H. sapiens* and *A. thaliana* genes are labeled, respectively.

For mammals, a very large proportion of all genes are syntenic and single copy (Figure 5a) as mentioned above. Smaller proportions of mammalian genomes are conserved and syntenic for duplicates or larger conserved multi-gene families. Interestingly, lineage-specific clusters were observed for most of the included mammalian clades. For example, we found lineage-specific clusters for Primates (such as the CENPJ example discussed above), Rodentia, Vespertilionidae, Felidae, Camelidae, and Bovidae (Figure 5a).

In contrast, in angiosperms only ~10% of clusters are syntenically conserved between eudicot and monocot species (Figure 5b). The remaining clusters are mostly lineage-specific clusters that appear as discrete columns (Figure 5b). This indicates that angiosperm genomes are highly fractioned and reshuffled, with abundant examples of specific clusters for particular phylogenetic lineages/plant families, such as Amaranthaceae, Brassicaceae, Poaceae, Fabaceae, Rosaceae, and Solanaceae (Figure 5b). Results also highlight species with more gene copies per cluster (e.g. orange/red rows), likely due to recent WGD events such as for *G. max*, *B. napus* and *P. trichocarpa* (Figure 5b).

Traditional phylogenetic profiling data typically show only the presence/absence of a gene family. Whereas, our synteny-based phylogenetic profiling is based on conserved genomic collinearity of gene families across lineages which provides potential novel information about changes of genomic context (transpositions and/or expansions) or the origin of “novel genes” of specific gene families. Such changes in genomic context provide intriguing candidate gene sets for investigating trait evolution.

### Synteny network for homeobox genes of mammals and angiosperms

To summarize and further illustrate synteny cluster properties between mammals and angiosperms species, we display synteny networks for the entire homeobox multi-gene family for both lineages (Figure 5c and 5d). For the mammals, the well-known Hox clusters, derived from WGD and tandem duplications [45, 46], were visualized as two huge clusters (*Hox1-8* and *Hox9-13*) connected by EVX gene cluster (*EVX1* and *EVX2*) (Figure 5c). ParaHox genes [47] *PDX1*, *GSX1*, and *GSX2* form one highly inter-connected cluster (Figure 5c), while the other three ParaHox genes *CDX1*, *CDX2*, and *CDX3* form respective independent clusters (Figure 5c). Moreover, we have found the synteny cluster of *DLX1-4*, and *DLX6* [48], cluster of *LHX2, 6*, and *9* [49], cluster of *NKX2-1* and *2-4* [50, 51], and cluster of *CERS5* and *6* [52] (Figure 5c).

Plant homeodomain proteins have been classified in the literature into various groups based on sequence similarity of their homeodomains [53–55]. Here the syntenic connections across the full set of homeobox genes provide novel insights to the origin and relationships of all homeobox subfamilies (Figure 5d). Some examples include conserved clusters (OCP3, RPL, and ATH1) [56–58]; WGD-derived clusters (*KNAT3-5,HAT1-3-HB2-HB4, HDG1-HDG7-ANL2-FWA,* and *HDG2-HDG3-PDF2-ATML1*) [59, 60]; eudicot-specific clusters (*STM, KNAT7, KNAT2-KNAT6, WOX1-PFS2* and *HB22-HB51*) [61–63], and monocot-specific clusters (i.e. *Os01g60270, Os06g04850, Os08g19590*) [64] (Figure 5d).

Synteny networks provide a complementary method to more traditional phylogenetic approaches for investigating the ancestry and homology relationships of (large) multi-gene families. For example, synteny information identified ancient tandem origins and lineage-specific transpositions of angiosperm MADS-box genes [22, 65, 66]. We have analyzed the mammalian homeobox genes. We clearly show and verify that the mammalian Hox genes appear as inter-connected synteny super-clusters and also find synteny connections to the ParaHox genes, consistent with the numerous previous reports [45–47]. In contrast, for plants we did not find any prominent tandem origin of homeobox clades, but did identify several examples of WGD-derived gene expansions and family-specific transpositions.

## Conclusions

Synteny analysis of multi-species genomics datasets has led to major advances in our understanding of evolutionary patterns and processes. However, few studies have systematically assessed and compared genomic properties across kingdoms [7]. Synteny network statistical parameters provide new possibilities for systematically evaluating gene (syntenic) diversification and/or conservation patterns over long evolutionary time scales. In this study, we have presented an analytic framework for large-scale synteny comparisons using network analysis of all suitable mammalian and angiosperm genomes. Assessment metrics based on synteny intuitively illustrate genome contiguity and copy number depth due to (paleo)polyploidy. The C-P method provides a means to characterize gene family dynamics in a comparative evolutionary context. We have displayed and compared features of all synteny clusters from these two important lineages and performed their clade-wide phylogenetic profiling. The results illustrate the dramatic differences in genomic dynamics within and between the two groups, exemplified by synteny networks of primate-specific gene transpositions (i.e. *CENPJ*), extant ohnologs surviving 2R of mammals, and for all mammal and angiosperm homeobox genes.

Dissection of the properties of all synteny clusters provides intriguing insights into the differing genomic architectures and dynamics of mammal and flowering plants. Examples in this study are just the tip of the iceberg. Much remains to be explored, but this study provides an intriguing foundation for future investigations to better understand genome evolution and elucidate regulatory mechanisms underlying diverse evolutionary biological processes. Such approach can further be extended to other phylogenetic groups and deeper evolutionary time scales.

## Methods

### Genome resources

All reference genomes were downloaded from public repositories (Supplemental Table 1). For each genome, we needed a FASTA format file containing peptide sequences of all predicted gene models, as well as a genome annotation file (GFF/BED) showing the positions of all the genes. Original gene names in the FASTA file have been modified into a prefix (unique identifier indicating species) and numeric GenBank gene ID. An in-house script was used for batch downloading genomes and modifying gene names.

All mammalian genomes were downloaded from NCBI. Initially we utilized the total list of available mammal genomes on NCBI (https://www.ncbi.nlm.nih.gov/genome/browse/). Using the list with our script, some records did not contain the complete required information for our analysis (i.e. no genome annotation files, or no FASTA file of total peptide sequences). In the end, we retrieved 87 mammalian genomes suitable for our analysis. Angiosperm genomes were collected from various public databases such as Phytozome (https://phytozome.jgi.doe.gov/pz/portal.html (Supplemental Table 1).)

### Peptide sequence annotation

For gene family annotation, we used HMMER (hmmscan) to perform domain annotations against the Pfam database (version downloaded: Pfam 30.0, Pfam-A with 16,306 entries) for all the peptides of the utilized genomes. Domains identified from one sequence were combined, and used for gene family annotation. Multiple occurrences of the identical domain within one protein were counted only once.

### Pairwise comparison, synteny blocks detection, and network construction

RAPSearch2 was used to perform all inter- and intra- pairwise all-vs-all protein similarity searchs. MCScanX was used for synteny block detection with default settings (window size: 50, number of match genes: >= 5). All outputting collinear files were integrated and curated into one tabular-format file, each row contains information about “Block_ID”, “Block_Score”, and syntenic gene pairs. This file creates a database which contains the entire syntenic nodes and syntenic connections derived from the input genomes. Detail procedures can be referred to a Github tutorial (https://github.com/zhaotao1987/SynNet-Pipeline).

### Network statistics

Network statistical analysis was carried out in the R environment (http://www.r-project.org), using the R package “igraph” [67]. We performed the analysis of the networks of mammal genomes and angiosperm genomes separately. The entire network must first be simplified to reduce duplicated edges (same syntenic pair may be derived from multiple detections), followed by the calculation of clustering coefficient, and node degree of each node.

We mapped gene family annotations to all the nodes, and computed the percentage for each gene family using its total occurrence in the synteny network against its total occurrence from the step “Peptide sequence Annotation“. We filtered gene families with at least 50 nodes and plot percentage against average clustering coefficient for all these gene families. Quartiles of percentage and average clustering coefficient was estimated according to their distributions. We describe values over Q3 (highest 25%) as high, and values below Q1 (lowest 25%) as low.

### Gene annotation enrichment analysis

Gene families of special interest (“high-high”, “high-low”, “low-high”, and “low-low”) were extracted from the total analysis. We then mapped gene(s) from the model species *H. sapiens* (for mammals) or *A. thaliana* (for angiosperms) to each of the gene families. We then performed online PANTHER overrepresentation test (http://pantherdb.org/) for each of the gene lists, with Bonferroni correction for multiple testing. In addition to the annotation of GO enrichment (biological process, molecular function, and celluar component), we also included analysis of “Reactome pathways”, “PANTHER pathways”, and “PANTHER protein class”. Results containing significant enriched terms was downloaded and illustrated as word clouds, by the R package “tagcloud”. Font sizes determined by “-log10(p-value)”. We depicted a maximum of the top 40 most significant terms.

### Network clustering and phylogenetic profiling

We used the infomap method to split the entire network, consisting of millions of nodes, into clusters [30]. Clustering results were determined by topological edge connections, edges were unweighted and undirected. All synteny clusters were decomposed into numbers of involved syntenic gene copies in each genome. Dissimilarity index of all clusters was calculated using the “Jaccard” method of the vegan package [68], then hierarchically clustered by “ward.D”, and visualized by “pheatmap”. We illustrate all the clusters of mammals (cluster size >= 2), and all angiosperm clusters with size >= 4.

## Figure Legends

**Supplementary Figure 1.**
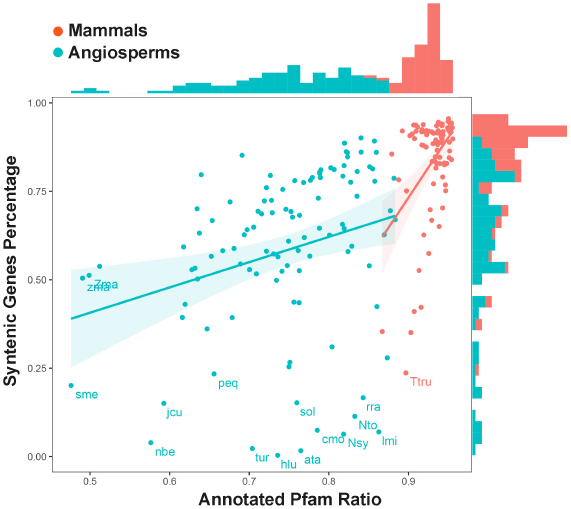
Plot of percentage syntenic genes again annotated (by Pfam) percentage of all genomes. Species were highlighted with abbreviated names if syntenic genes percentage lower than 0.25 or annotated proteins (by Pfam) lower than 0.5.

**Supplementary Figure 2.**
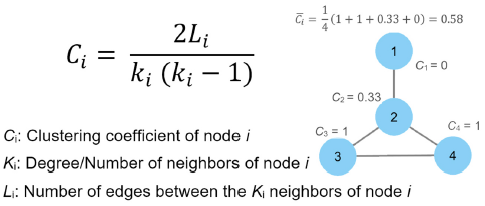
Schematic diagram for the calculation of the average clustering coefficient.

**Supplementary Table 1** Mammalian and angiosperm genomes used in this study

**Supplemental Table 2** Gene families with significant C-P features of mammals and angiosperms.

## Declarations

Ethics approval and consent to participate

Not applicable.

## Availability of data and material

Data-sets and computer code used in this study are available at DataVerse: (https://dataverse.harvard.edu/privateurl.xhtml?token=308d70cc-f489-435d-b7a5-f4fc5acd4842). This includes the modified FASTA and BED files of all mammal and angiosperm reference genomes. The scripts for network database preparation (pairwise comparison, synteny block detection, and data integration), Pfam domain annotation, network clustering and statistics, phylogenetic profiling, and for the figure preparation (if applicable) are all included.

## Authors’ contributions

TZ and MES designed the study, TZ assembled the genomic data and performed the analysis. TZ and MES wrote the paper. All authors read and approved the final manuscript.

## Acknowledgements

TZ was supported by China Scholarship Council. Symbols for diagrams courtesy of the Integration and Application Network (ian.umces.edu/symbols).

